# h5adify: neuro-symbolic metadata harmonization enables scalable AnnData integration with local large language models

**DOI:** 10.64898/2026.02.28.708740

**Authors:** Lucas Rincón de la Rosa, Abdelmalek Mouazer, Masoud Navidi, Edgar Degroodt, Tamara Künzle, Sylvain Geny, Ahmed Idbaih, Maïté Verrault, Karim Labreche, Isaïas Hernández-Verdin, Agustí Alentorn

## Abstract

**Background:** The rapid growth of public single-cell and spatial transcriptomics repositories has shifted the main bottleneck for atlas-scale integration from data generation to metadata heterogeneity. Even when datasets are released in the AnnData H5AD format, inconsistent column naming, partial annotations, and mixed gene identifier conventions frequently prevent reproducible merging, downstream benchmarking, and reuse in foundation model training. Automated approaches that resolve semantic inconsistency while preserving biological validity are therefore essential for scalable data reuse.

**Results:** We present `h5adify`, a neuro-symbolic toolkit that combines deterministic biological inference with locally deployed large language models to transform heterogeneous AnnData objects into schema-normalized, integration-ready representations. The framework performs metadata field discovery, gene identifier harmonization, optional paper-aware extraction, and consensus resolution with explicit uncertainty logging. Benchmarking four open-weight model families deployed through Ollama (Gemma, Llama, Mistral, and Qwen) demonstrates that small local models achieve high semantic accuracy in metadata resolution with low hallucination rates and modest computational requirements. In controlled simulations introducing annotation noise into single-cell and Visium-like datasets, harmonization improves integration benchmarking and reduces spurious batch effects. Application to sex-annotated glioblastoma datasets recovers biologically coherent microenvironmental patterns and cell type–specific genomic differences not explained by differential expression alone.

**Conclusions:** Together, `h5adify` provides a reproducible framework for evaluating LLM-assisted biocuration and enables scalable, privacy-preserving metadata harmonization for modern single-cell atlases and foundation model pipelines. These results demonstrate that modular neuro-symbolic integration of deterministic biological inference and small local language models can effectively resolve semantic heterogeneity while remaining computationally accessible.

## 1 Introduction

Single-cell RNA sequencing and spatial transcriptomics have become central technologies for dissecting tissue composition, cellular states, and microenvironmental organization in development and disease. This growth has been accompanied by a rapid expansion of public repositories and atlas initiatives, which increasingly distribute processed matrices in the AnnData H5AD format [1] and are commonly analyzed through the Scanpy and Squidpy ecosystems [2, 3]. Parallel infrastructure efforts, including CZ CELLxGENE, have improved access and interactive exploration of curated datasets [4].

However, atlas scale reuse is still impeded by a less visible layer of heterogeneity: dataset metadata. In practice, different studies encode the same concepts using divergent column names, levels of granularity, and local conventions. Examples include the inconsistent use of `donor`, `patient`, `sample` and `batch` fields, or the presence of free-text disease labels that mix histology, grade, molecular subtype, and treatment context. These inconsistencies create failure modes that are difficult to detect downstream, such as spurious batch effect metrics driven by mislabeled covariates, incorrect stratification in sex-aware analyses, or silent loss of samples during merges because of ambiguous identifiers.

This issue has become more pressing with the emergence of single-cell foundation models and transfer learning pipelines, which depend on large corpora of harmonized data. Recent models pre-trained on tens of millions of cells, such as Geneformer [5],scGPT [6], and scFoundation [7], highlight the value of large-scale pretraining for representation learning and downstream prediction. Yet the reliability of such models is constrained by the quality of the input corpus: metadata heterogeneity can introduce label leakage, systematic confounding, or unwanted domain shifts that remain unquantified. In addition, model development often prioritizes expression normalization and batch correction while the upstream biocuration steps that define *what* is being integrated are rarely evaluated explicitly [8].

Several resources address related challenges in bulk and functional genomics. MetaSRA normalizes Sequence Read Archive sample metadata by mapping to controlled terms [9], recount3 provides uniformly processed RNA sequencing data with standardized access layers [10], and GEOMetaCuration supports collaborative manual curation of GEO studies [11]. In the single-cell field, the community has focused heavily on computational integration methods and benchmarking, including scVI and scANVI [12, 13] and multi-method integration frameworks [14–16]. Yet, even the best integration methods cannot compensate for missing or inconsistent metadata fields, and few tools provide a systematic way to discover, resolve, and audit metadata mappings across heterogeneous H5AD files.

Here, we introduce `h5adify`, a neuro-symbolic metadata harmonization toolkit designed to make metadata heterogeneity *measurable* and *actionable* in AnnData based workflows. The key idea is to treat metadata harmonization as a consensus problem: deterministic biological signals (gene naming patterns, sex chromosome expression, dataset structural cues) are combined with semantic reasoning provided by local large language models, optionally grounded in the associated publication. This design yields a modular pipeline that can operate in offline environments through Ollama, and produces explicit logs that support reproducibility, failure analysis, and benchmarking of curation quality. We evaluate `h5adify` through multi-dataset benchmarks, controlled simulations, and two glioblastoma case studies (single-cell and spatial) that illustrate how harmonization enables sex-aware analyses beyond differential expression, including spatial organization and inferred intercellular communication.

## 2 Results

### 2.1 Local large language models achieve high semantic accuracy for metadata harmonization

We first quantified the accuracy and efficiency of metadata harmonization under four local model families deployed through Ollama, using four benchmark datasets with heterogeneous annotation conventions (Tasic et al., Han et al., Almanzar et al., and Travaglini et al.) [17–20]. Across datasets, field resolution accuracy was consistently high for structurally constrained fields (donor, sex, technology) and remained robust for more ambiguous fields (batch and disease), where semantic reasoning is required to map free-text descriptions to standardized schema fields.

Figure 1 summarizes these results in a single multi-panel view. The benchmark reveals that model performance differences are driven largely by ambiguous metadata organization rather than by biological inference. Importantly, the evaluation highlights the complementarity of exact and semantic matching: strict column-name agreement underestimates practical usability, because datasets frequently encode the correct variable under alternative labels, while value-level concordance captures the intended mapping. Runtime profiling further demonstrates that local execution is feasible without high-end accelerators, supporting deployment in privacy constrained environments where data cannot be shared externally.

**Fig. 1:**
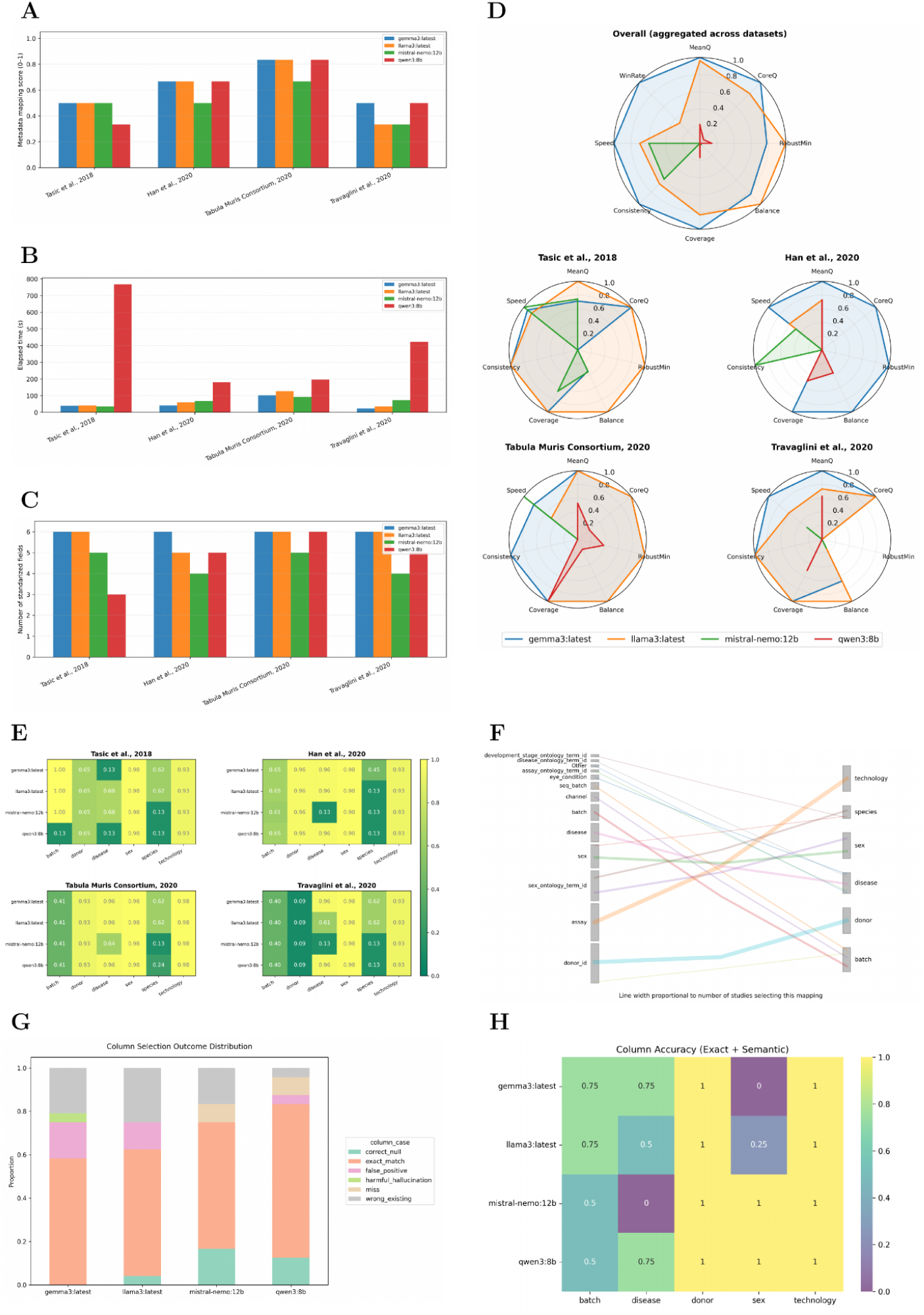
Benchmarking local large language models for metadata harmonization. (A) Normalized mapping score per study. (B) End-to-end runtime, including prompt execution and parsing. (C) Number of standardized fields produced per study. (D) Radar summaries of accuracy, robustness and coverage, shown overall and per benchmark dataset. (E) Field-level exact-match heatmaps, where darker values indicate closer agreement to gold standards. (F) Sankey overview of common column-to-field mappings learned by each model family, highlighting convergence on a shared vocabulary. (G) Distribution of mapping outcomes, including correct, exact-match, semantic-match and error categories. (H) Field-wise accuracy aggregated across datasets.

### 2.2 Controlled simulations show that metadata harmonization improves integration benchmarking

We next evaluated the impact of metadata harmonization on downstream integration benchmarking using controlled simulations that inject annotation noise, inconsistent naming conventions, and structured batch effects in both single-cell and Visium-like spatial datasets (Methods). These simulations mimic common real-world failure modes, including incomplete sex labels, ambiguous donor identifiers, and technology fields encoded inconsistently across studies.

Figure 2 reports scIB benchmarking results before and after `h5adify` harmonization across three simulation scenarios. In the single-cell settings, harmonization improves the interpretability of integration metrics by ensuring that batch and donor variables correspond to comparable biological entities across simulated studies. In the spatial setting, where spot-level metadata and coordinate systems amplify the consequences of inconsistent identifiers, harmonization reduces spurious metric variability and yields more stable comparisons across integration approaches. These results emphasize that integration benchmarks depend not only on the correction method, but also on the quality of the metadata defining the benchmark axes.

**Fig. 2:**
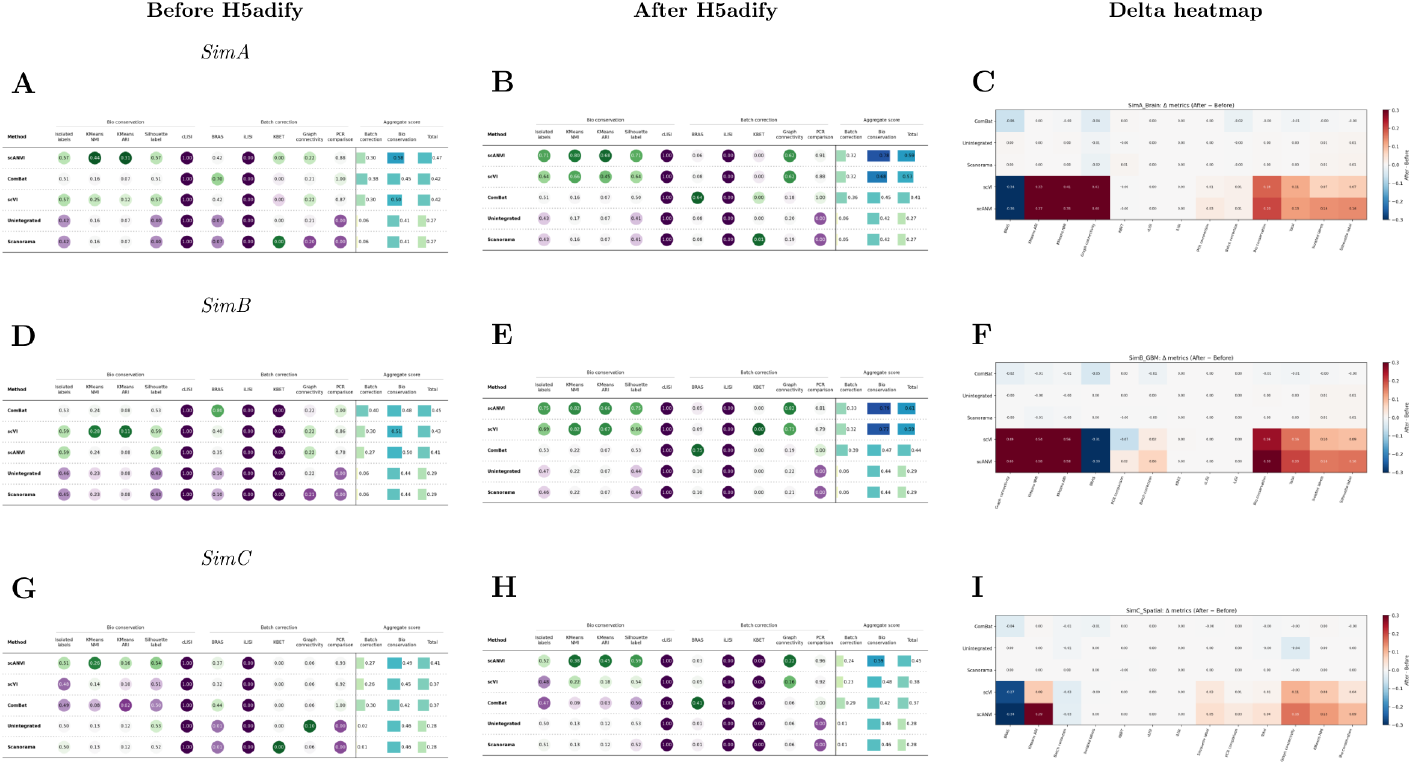
Simulation benchmarking of integration before and after metadata harmonization. SimA (A–C) models a human brain single-cell setting with structured donor and batch effects; SimB (D–F) models a glioblastoma-like single-cell setting with stronger confounding; SimC (G–I) models a Visium-like spatial setting with multi-factor effects (technology, tissue section, and donor) and noisy metadata. For each scenario, we report scIB metric tables before (A,D,G) and after (B,E,H) harmonization, and the corresponding metric deltas (C,F,I).

As shown in Figure A1 we achieved a virtually perfect classification of sex and species in the different simulations (either using scRNA or spatially resolved transcriptomics simulated data).

### 2.3 Sex-associated differences in glioblastoma emerge beyond differential expression in single-cell data

We then investigated whether harmonization enables sex-aware biological analysis in glioblastoma, focusing on single-cell data comprising malignant, immune, and vascular compartments. Figure 3 summarizes the main findings. The global embedding shows that sex labels are broadly intermingled across cell states (Fig. 3A), while cell type specific stratification remains the primary driver of transcriptional structure (Fig. 3B). Consistent with this, the distribution of `XIST` expression aligns with sex labels and supports internal quality control of the metadata harmonization outcome (Fig. 3D).

**Fig. 3:**
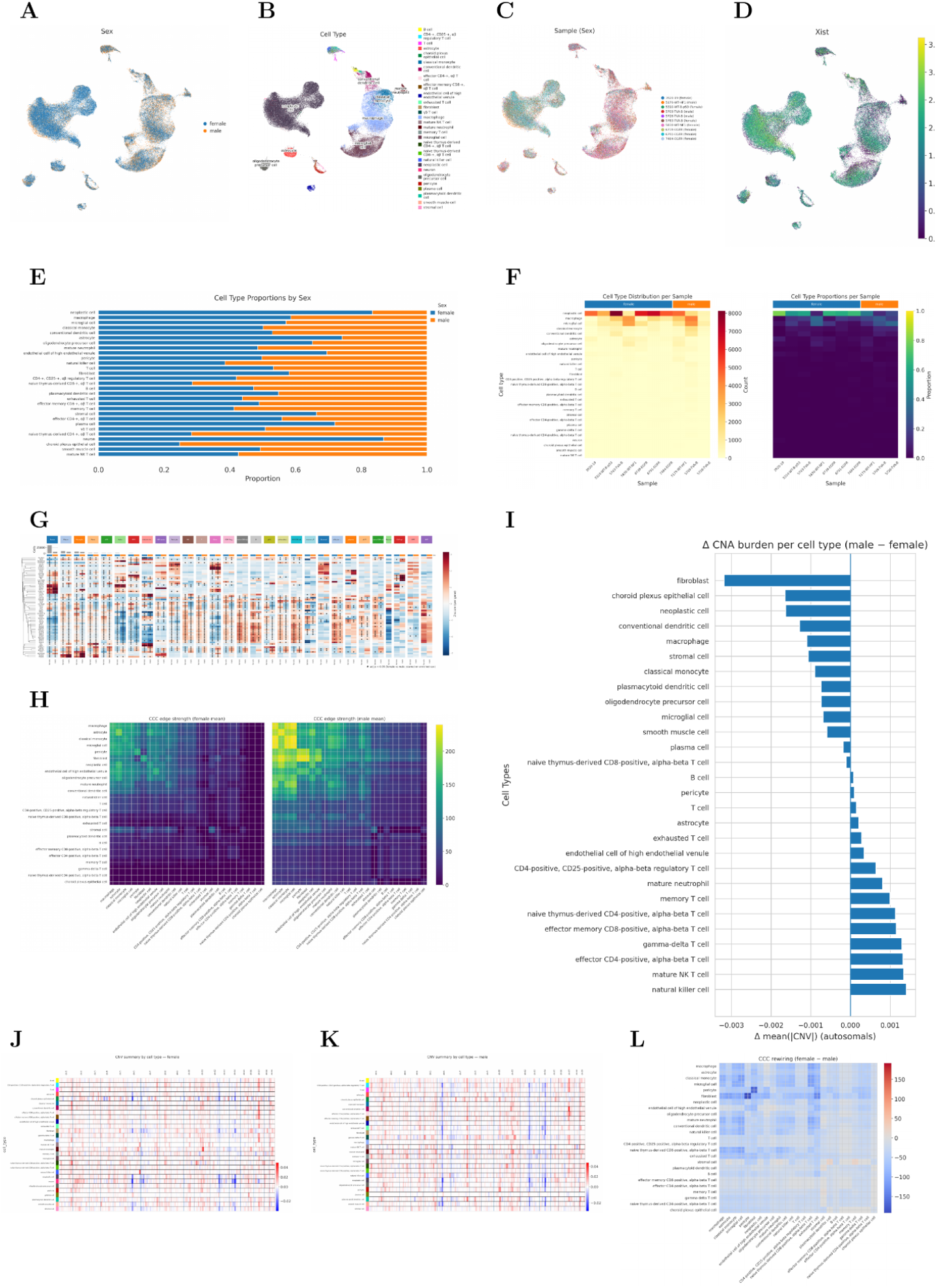
Single-cell glioblastoma analysis stratified by sex (Soni et al.). (A) UMAP colored by sex. (B) UMAP colored by annotated cell type. (C) UMAP colored by sample (sex-coded). (D) XIST expression on the embedding as an internal control for sex annotation. (E) Cell-type proportions by sex. (F) Cell-type distributions by sample shown as counts (left) and proportions (right). (G) Genome-wide inferred CNV heatmap across cells. (H) Cell–cell communication edge-strength matrices (female and male means). (I) Mean autosomal CNV burden difference per cell type (male minus female). (J,K) CNV summaries by cell type for female and male, respectively. (L) Difference heatmap of inferred interaction strengths (female minus male).

Despite limited separation in expression space, downstream analyses reveal sex-associated differences in genomic instability and inferred intercellular programs. Copy number burden summaries indicate that multiple non-malignant compartments, including pericytes and smooth muscle like cells, exhibit sex-associated shifts in inferred autosomal copy number deviation (Fig. 3I–K). In parallel, cell–cell communication analysis identifies sex-associated rewiring patterns across immune and vascular interactions (Fig. 3H,L). Notably, these signals are not mirrored by a uniformly large number of differentially expressed genes, as supported by the extensive set of cell-type specific volcano plots (Fig. 4), suggesting that sex-associated differences can manifest through coordinated pathway and interaction changes rather than through strong single-gene effects.

**Fig. 4:**
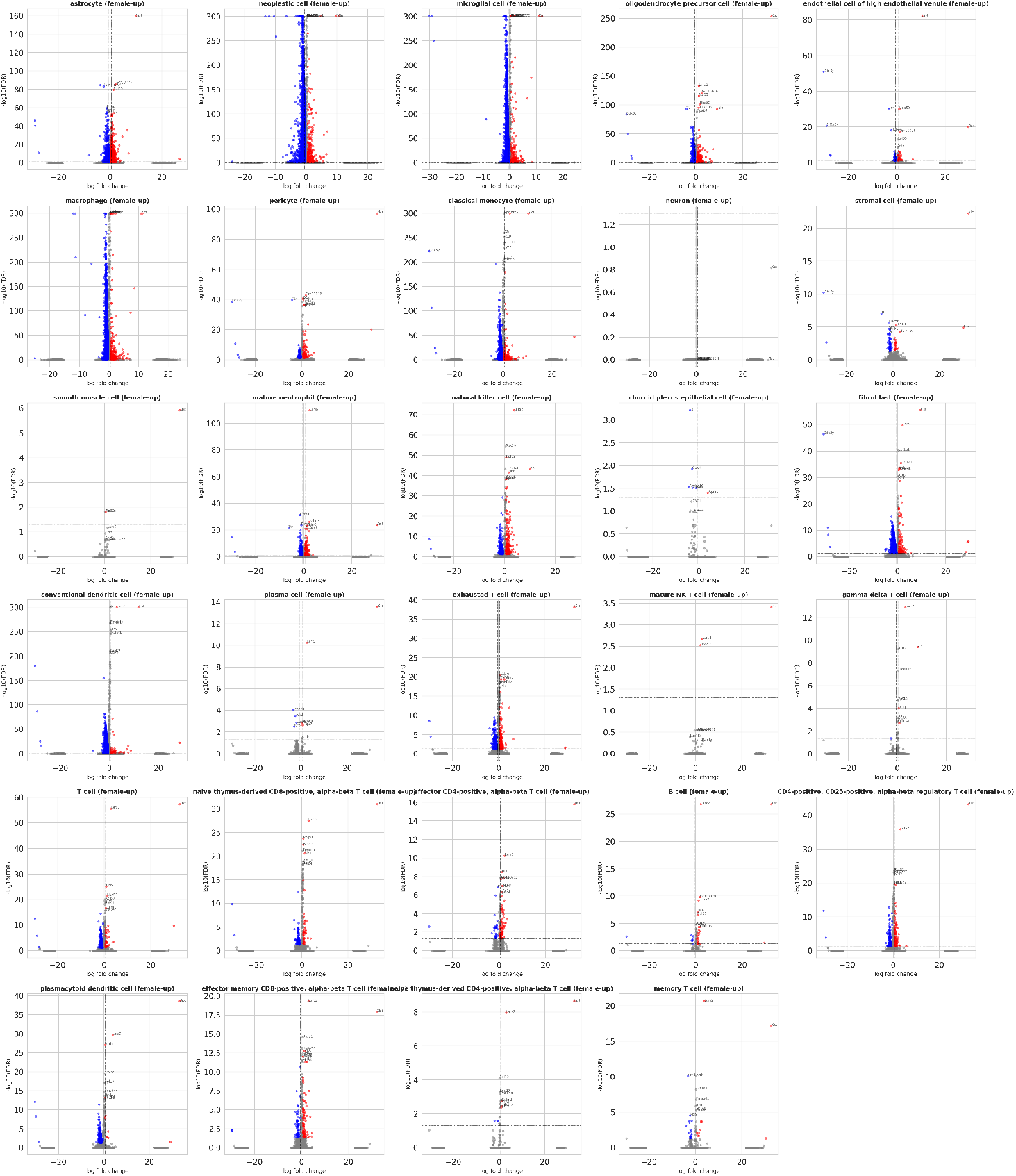
Supplementary differential expression summary. Volcano plots for multiple cell types comparing female versus male within each compartment. The limited number of strong effects in several cell types contrasts with the broader sex-associated differences observed in copy number summaries and inferred intercellular communication, supporting the interpretation that sex effects can be mediated by coordinated multivariate programs.

### 2.4 Spatial hubs and microglia organization differ by sex in glioblastoma IDH-wildtype

Finally, we analyzed a Visium spatial transcriptomics cohort restricted to glioblastoma IDH-wildtype samples (Methods) and used Starfysh to infer spot-level cell type compositions and discover spatial hubs [21]. Figure 5 summarizes the spatial pipeline and sex-stratified results. The embedding and hub decomposition reveal reproducible spatial programs across samples, and hub abundance summaries suggest that a subset of hubs exhibit sex-associated shifts in prevalence across patients.

**Fig. 5:**
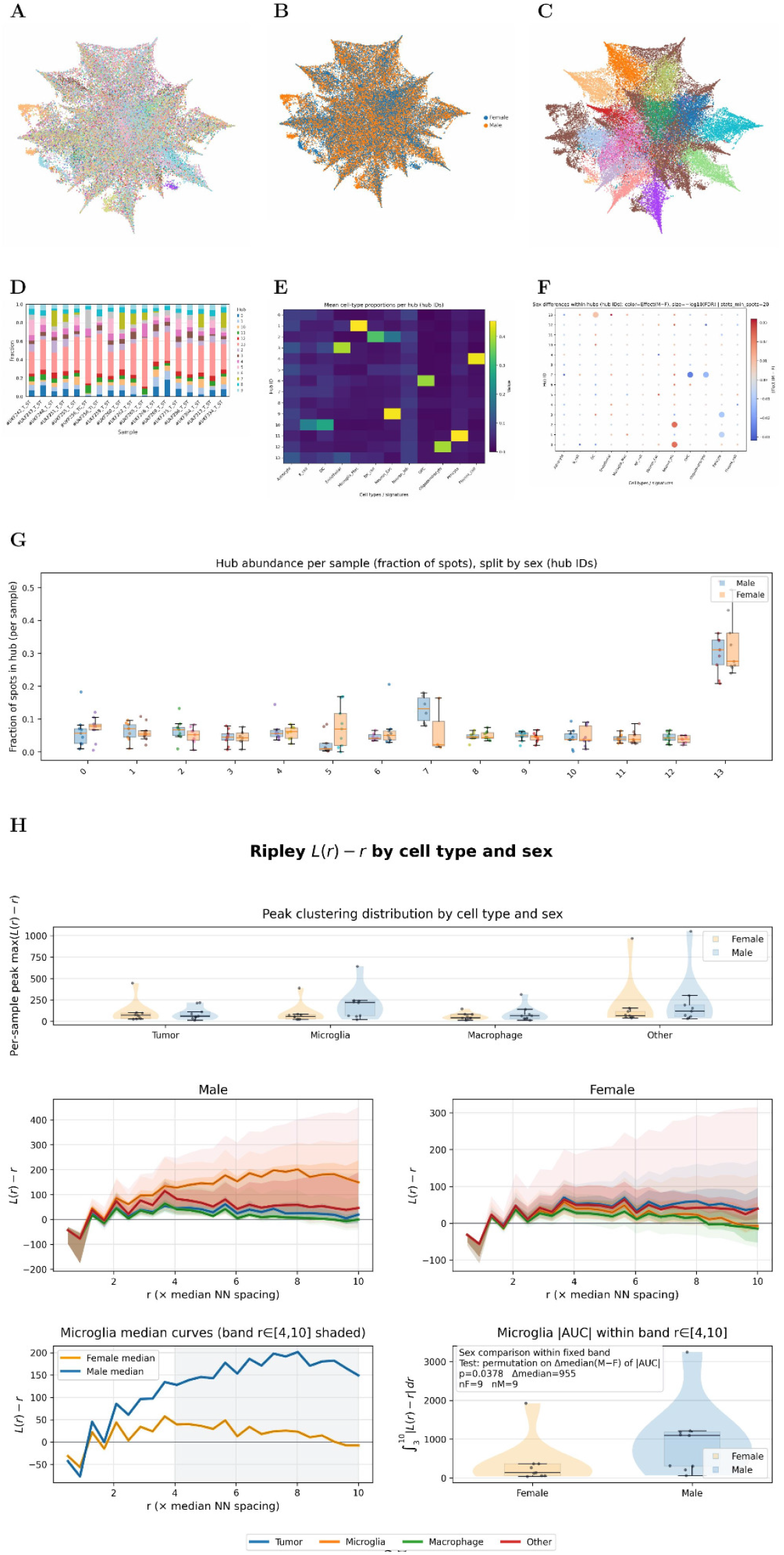
Spatial transcriptomics analysis of glioblastoma IDH-wildtype stratified by sex (Ravi et al. [22]). (A) UMAP of all spots. (B) UMAP colored by sex. (C) UMAP colored by Starfysh-derived hubs. (D) Hub composition per sample. (E) Mean cell-type proportions per hub. (F) Sex-differential hub enrichment (dot size encodes effect size; color encodes direction). (G) Hub abundance per sample split by sex. (H) Ripley *L*(*r*) − *r* analysis by cell type and sex, including peak clustering distributions and microglia-focused band statistics.

To quantify sex-associated differences in the spatial organization of specific compartments, we applied Ripley-based spatial statistics to compare clustering profiles between male and female samples. The strongest signal was observed in microglia, for which we identify a radius band of consistent departure from complete spatial randomness and observe a significant difference in the magnitude of clustering within this band between sexes (Fig. 5H). This result supports the interpretation that sex-associated glioblastoma dimorphism can manifest not only in immune cell composition, but also in the spatial arrangement of immune niches and their proximity to tumor-associated hubs.

## 3 Discussion

### 3.1 Metadata harmonization as a prerequisite for atlas-scale reuse and foundation model training

The field has reached a stage where the availability of public single-cell and spatial data is no longer the limiting factor for integrative analyses. Instead, metadata heterogeneity has become a primary barrier to reuse. This barrier is subtle because it often does not surface as an explicit error: rather, it propagates into downstream analyses as silent confounding, inconsistent stratification, and irreproducible merges. The problem is particularly acute for foundation model pipelines, where training corpora aggregate diverse studies and where systematic annotation mismatches can become encoded as spurious structure in learned representations [5–7].

`h5adify` addresses this gap by treating metadata harmonization as a first-class, testable task, rather than as a manual preprocessing step. By combining deterministic biological evidence with semantic reasoning from local large language models, the pipeline creates an auditable chain of decisions that can be benchmarked and compared across model variants. Our simulation results further show that harmonization is not only a curation convenience: it directly affects the stability and interpretability of integration benchmarks. These findings align with broader benchmarking work emphasizing that end-to-end pipelines are only as reliable as the assumptions used to define batches, donors, and biological strata [8, 23].

### 3.2 Positioning relative to existing tools

Several existing resources address aspects of metadata normalization, but they typically operate outside the AnnData ecosystem or focus on bulk data. MetaSRA provides ontology-guided normalization for Sequence Read Archive samples [9], recount3 focuses on uniformly processed bulk RNA sequencing at scale [10], and GEOMetaCuration supports collaborative manual curation for GEO studies [11]. In the single-cell domain, integration methods such as Seurat, Harmony, Scanorama, and scVI target expression-level alignment [12–16], while platforms such as CZ CELLxGENE prioritize standardized access and interactive exploration [4].

`h5adify` is complementary to these efforts. Rather than replacing existing integration methods, it focuses on upstream harmonization and auditability within the AnnData representation. This emphasis is important because integration methods assume that key covariates have been defined correctly, yet public datasets frequently encode these covariates in inconsistent ways. In addition, recent LLM-based systems have begun to address metadata curation using agentic workflows and structured retrieval [24, 25]. Related efforts include agentic ingestion and standardization frameworks [26], chat-based exploration of single-cell data [27], and LLM-assisted automated annotation and prior-informed multi-dataset integration [28]. Our work extends this direction by explicitly combining LLM reasoning with biological inference and by targeting the practical interface where researchers manipulate data objects: the H5AD files used in analysis and in model training pipelines.

### 3.3 Local LLMs enable privacy-preserving, resource-efficient biocuration

A practical constraint in biomedical settings is that data cannot always be shared with external services. Local inference through Ollama provides a pragmatic solution, enabling the use of open models for semantic normalization while keeping raw data on site. Our benchmark indicates that compact models can achieve high semantic accuracy for core metadata fields, and that their performance is largely limited by ambiguity in the dataset annotations rather than by a lack of biological signal. Importantly, runtime and memory profiles are compatible with CPU execution or consumer GPUs, lowering the barrier for routine adoption in academic and clinical environments.

### 3.4 Sex-associated glioblastoma signals extend beyond differential expression

The glioblastoma case studies illustrate why metadata harmonization matters for downstream biological interpretation. Sex is often inconsistently encoded, and analyses that stratify by sex or gender can fail silently when labels are incomplete or contradictory. After harmonization and internal quality control using sex-chromosome markers, we identify sex-associated differences that extend beyond simple differential expression and are consistent with the increasing use of expression-derived CNV surrogates to characterize non-malignant programmes in tumor ecosystems [29]. In single-cell data, inferred copy number burden differences in pericytes and smooth muscle-like cells suggest sex-associated variation in vascular and perivascular programs, consistent with evidence that neurovascular and blood–brain barrier regulation can be sex dependent, including sex- and age-associated differences in pericyte-related gene regulation [30] and reference atlases describing molecular diversity across human brain vascular and perivascular compartments [31]. Moreover, the observed cell–cell communication rewiring indicates that sex-associated differences can manifest as changes in coordinated interaction programs, which may be weak at the single-gene level but pronounced when aggregated across ligand–receptor systems [32–35].

In spatial data restricted to glioblastoma IDH-wildtype samples, our results support sex-associated differences in the organization of myeloid compartments, particularly microglia clustering patterns. This observation is concordant with recent reports of sexually dimorphic myeloid biology in glioblastoma and related glioma contexts, including sex-specific roles for suppressive myeloid subsets and differences in microglia and macrophage states [36–39]. The spatial framing is important: differences in infiltration can coexist with differences in spatial arrangement, and niche organization may better reflect functional immune programs than cell proportions alone. More generally, recent spatial atlases of glioblastoma highlight that both malignant and non-malignant programs are strongly spatially structured [22, 40]. Our results suggest that sex can modulate this spatial structure in the non-neoplastic compartment, providing a testable hypothesis for future prospective studies.

### 3.5 Limitations and future directions

This work has several limitations. First, although we provide both simulation and real-data validations, the space of metadata heterogeneity is effectively unbounded and is shaped by evolving community conventions. Second, sex-associated analyses are sensitive to sampling, treatment context, and clinical confounders; therefore, our biological conclusions should be interpreted as hypotheses supported by multiple analysis modalities rather than as definitive causal claims. Third, large language models remain probabilistic systems; while our consensus layer reduces hallucination risk, careful auditing and transparent reporting remain essential.

Future work will focus on extending the target schema toward richer ontologies, improving provenance tracking for harmonized fields, and integrating automated tests that detect downstream inconsistencies early (for example, incompatible batch variables or implausible sex assignments). In parallel, scaling harmonization to multimodal datasets that couple transcriptomics with histology, imaging, and clinical covariates will be critical for next-generation digital twin and foundation model efforts.

## 4 Conclusions

`h5adify` introduces a reproducible, neuro-symbolic approach to metadata harmonization for AnnData objects that combines deterministic biological inference with local large language models. Across benchmarks and controlled simulations, we show that harmonization improves the stability and interpretability of integration analyses and enables robust reuse of heterogeneous datasets. In glioblastoma case studies, harmonization supports sex-aware analyses that recover differences in spatial myeloid organization, inferred copy number burden, and interaction programs that are not captured by differential expression alone.

By providing explicit audit logs and supporting fully local execution, `h5adify` lowers the barrier for privacy-preserving biocuration and creates a practical bridge between the growing ecosystem of public single-cell resources and the data requirements of atlas-scale integration and foundation model training.

## 5 Methods

### 5.1 Software overview and design principles

`h5adify` is implemented in Python and targets AnnData objects as the primary data representation [1]. The pipeline follows a neuro-symbolic design in which deterministic rules and biological constraints provide high precision anchors, while local large language models contribute flexible semantic reasoning for heterogeneous metadata (Fig. 6). The implementation supports command-line usage and a graphical interface, and logs all intermediate decisions (including confidence scores, candidate mappings, and conflict resolution outcomes) to facilitate reproducibility and error analysis.

**Fig. 6:**
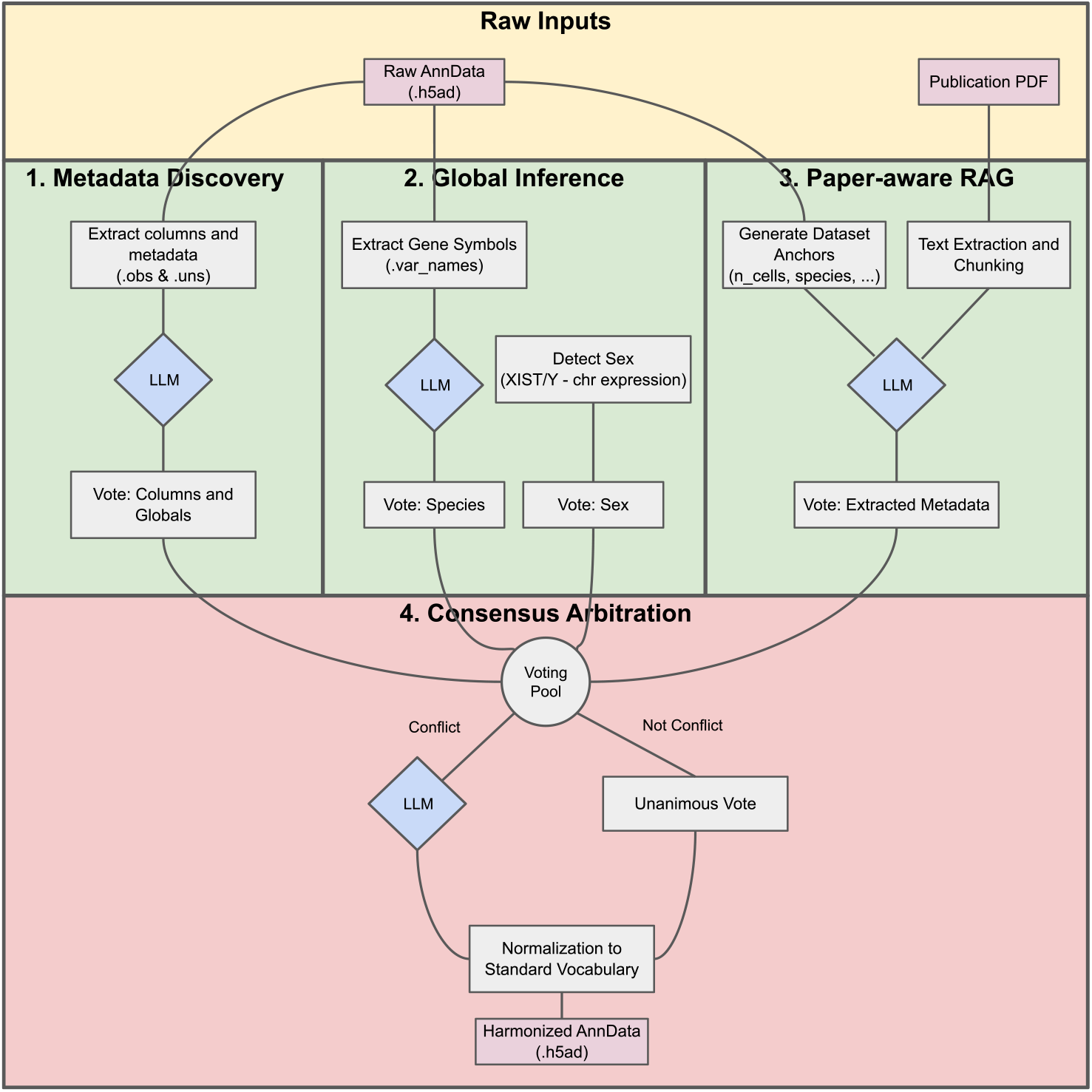
Overview of the `h5adify` pipeline. Metadata evidence is gathered from three complementary modules: (i) structural metadata extraction from `.obs` and `.uns`, (ii) deterministic biological inference from expression and gene identifier patterns, and (iii) optional paper-aware extraction from the associated publication. Evidence is combined through an explicit voting pool and conflict resolution layer, yielding a normalized AnnData object with standardized vocabulary and audit logs.

### 5.2 Deterministic biological inference

#### Gene identifier type and species

Gene identifiers were harmonized using canonical references obtained from the Ensembl database. For each dataset, gene symbols in `adata.var` were mapped to standardized identifiers (e.g., HUGO gene nomenclature for human datasets) via online Ensembl queries. Original identifiers were preserved in an auxiliary column to ensure reversibility and traceability of transformations.

#### Sex inference

Biological sex was inferred directly from gene expression using a deterministic scoring scheme based on aggregate Y-chromosome expression (e.g., *DDX3Y, KDM5D, UTY* ) and X-inactivation markers (*XIST* ). For each observation *i*, we computed:

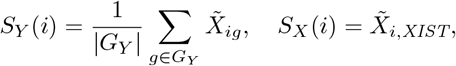

where *G*_*Y*_ denotes a curated set of Y-linked genes and 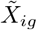 represents library-size normalized log-expression for gene *g* in observation *i*. For each observation, we computed a continuous delta score defined as the difference between normalized Y-linked expression and XIST expression. This score was thresholded to produce categorical predictions (male, female, unknown).

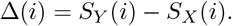

Samples were classified as:

- male if Δ(*i*) *> τ*_*m*_,
- female if Δ(*i*) *<* −*τ*_*f*_,
- unknown otherwise,

where thresholds *τ*_*m*_ and *τ*_*f*_ were selected via grid search to maximize balanced accuracy on benchmark datasets.

In spatial transcriptomics datasets, sex marker expression may be attenuated due to reduced transcript capture sensitivity and spot-level cellular heterogeneity, potentially limiting separability relative to dissociated single-cell data.

### 5.3 Local large language models and prompt structure

We deploy open weight instruction tuned models through Ollama, enabling fully local inference without data exfiltration. The benchmarked models correspond to Gemma, Llama, Mistral and Qwen families (Fig. 1) and were selected for their availability, strong general reasoning performance, and compatibility with CPU execution or consumer GPUs (less than 16 GB of memory in our tested configurations).

Prompts are task specific and decomposed into three roles. (i) An *Indexer* prompt enumerates candidate metadata fields and proposes mappings from dataset specific columns to a target schema. (ii) A *Researcher* prompt optionally incorporates signals extracted from the associated publication. (iii) An *Arbiter* prompt is invoked when votes conflict and produces a final decision together with a short justification that is stored in the audit log.

### 5.4 Benchmark datasets and evaluation protocol

To evaluate `h5adify`’s performance, we curated a set of publicly available single-cell transcriptomics datasets with diverse biological contexts, technologies, and species representation. All datasets were accessed via CellxGene or related portals and are associated with peer-reviewed publications: Tasic et al. (mouse cortex), Han et al. (human cell landscape), Almanzar et al. (Tabula Muris), and Travaglini et al. (human lung) [17–20].

Gold-standard annotations were generated by a single expert curator through deterministic inspection of AnnData objects and associated publications. Because target fields correspond to explicitly documented technical or clinical descriptors, subjectivity was limited relative to interpretative biological annotations. When multiple plausible columns existed (e.g., human-readable versus codified encodings such as `sex` versus `sex id`), the human-readable column was selected as the gold standard.

#### Column discovery outcomes

Column selection accuracy was defined as the proportion of predicted source columns that matched the gold-standard annotation. Predictions were considered correct if they either:

1. Matched the exact column name, or
2. Exhibited high semantic concordance with the ground truth (Adjusted Rand Index ≥ 0.9). This accounts for cases in which the model selects a codified or encoded representation of the correct field (e.g., numeric sex codes instead of human-readable labels), preserving semantic equivalence while distinguishing structural preference.

The threshold of 0.9 was selected to require near-identical categorical concordance while tolerating differences in encoding format.

Semantic concordance was quantified using the Adjusted Rand Index (ARI):

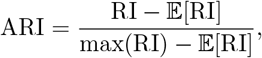

where RI denotes the Rand Index and 𝔼 [RI] its expected value under random assignment.

Null detection accuracy was defined as the proportion of cases where the model correctly returned “none” when no valid source column existed.

#### Field resolution accuracy

For fields represented by explicit columns, resolution accuracy corresponds to the successful selection of the correct column (exact or semantic match). For global fields (for example, species), accuracy corresponds to the inferred value. We additionally report runtime and a set of aggregate quality indicators summarizing the stability of model using precision, recall, and F1-score relative to manually curated annotations.

#### Hallucination Rate

Operationally, hallucinations were identified when the predicted column or value was neither present in the AnnData object nor recoverable from publication text or biological inference modules. We distinguished:

- Benign hallucinations: Incorrect but biologically plausible outputs.
- Harmful hallucinations: Outputs contradicting molecular evidence or publication content.

Hallucination rate was calculated as the proportion of unsupported predictions across all evaluated fields.

### 5.5 Simulation framework for controlled metadata heterogeneity

To quantify the impact of metadata harmonization in a controlled setting, we implemented a simulation suite that generates single-cell and Visium-like spatial datasets with injected annotation noise and inconsistent naming conventions. Single-cell simulations sample cell type compositions and generate counts using a negative binomial model with cell type specific mean profiles. We introduce donor-specific effects by multiplicative perturbations of gene means, and technology-specific effects by perturbing a subset of genes with stronger variance.

Spatial simulations use Spider [41] as a generative scaffold for Visium-like coordinates and cell-type mixture fields. Briefly, Spider defines spatial score surfaces *s*_*k*_(*x, y*) for each cell type *k* by sampling spatial patterns that include cold (localized), mixed (weakly structured) and compartmentalized (strongly segregated) regimes through a cell-type transition matrix and optional continuous gradients.

Scores are exponentiated and normalized to probabilities, *p*_*k*_(*x, y*) = exp(*s*_*k*_(*x, y*) − max_*j*_ *s*_*j*_(*x, y*))*/* ∑_*j*_ exp(*s*_*j*_(*x, y*) − max_*j*_ *s*_*j*_(*x, y*)), and spot-level mixtures are sampled from *p*_*k*_(*x, y*) before generating counts with a negative binomial model. In SimC, we additionally simulate multi-factor structure by assigning each spot to a donor, tissue section, and technology, and we apply factor-specific perturbations to a subset of genes to emulate structured unwanted variation. To reproduce realistic harmonization failure modes, we inject heterogeneous and partially missing metadata columns (for example, inconsistent sample identifiers, alternative donor encodings, multiple batch-like fields, and incomplete sex annotation). These simulated datasets are then used to benchmark downstream integration methods before and after harmonization.

### 5.6 Integration benchmarking with scIB

We assess integration quality using the scIB benchmarking framework [23]. For each simulated dataset, we compare representative integration strategies spanning classical correction and latent variable models: unintegrated analysis, ComBat [42], Scanorama [16], scVI and scANVI [12, 13]. Metrics include measures of biological conservation and batch mixing (for example, silhouette-based scores and neighborhood mixing statistics), reported in the simulation figures.

### 5.7 Glioblastoma case studies

#### Single-cell RNA sequencing analysis

We analyzed a sex annotated glioblastoma single-cell dataset with malignant, immune, and vascular compartments. Cells were filtered using standard quality control criteria, normalized, and embedded using PCA and UMAP as implemented in Scanpy [2]. Cell types were defined by the reference annotations provided by the original study and harmonized through `h5adify`. Differential expression by sex was performed within each cell type using log fold change thresholds and multiple testing correction.

Copy number alterations were inferred from expression by comparing smoothed gene expression along genomic coordinates and aggregating deviations into per-cell and per-cell-type burden scores, following standard principles used in scRNA-seq CNA inference tools such as inferCNV and CopyKAT [29]. We report autosomal CNV summaries to reduce sex chromosome driven artefacts and emphasize between-sex differences within each annotated compartment.

Intercellular communication was characterized through ligand–receptor interaction inference and summarized as directed edge-strength matrices between sender and receiver cell types. We used curated ligand–receptor resources and statistical testing strategies aligned with CellPhoneDB and CellChat, and we complemented pairwise interaction summaries with downstream pathway-centric interpretation [32–35].

#### Spatial transcriptomics analysis and sex stratification

For spatial analyses we focused on glioblastoma IDH-wildtype samples from a published Visium cohort [22, 40]. Spatial deconvolution and hub discovery were performed using Starfysh [21] on a per-sample basis. We then merged the per-sample results onto a unified backbone AnnData object, preserving H5AD-safe summaries and storing detailed run reports as external JSON artefacts.

#### Spatial hub statistics

Hub abundance per sample was computed as the fraction of spots assigned to each hub. Sex differences were assessed using non-parametric tests across samples, with visual summaries reported as boxplots.

#### Ripley’s L for sex-associated clustering

We used Ripley’s K function and its stabilized L transform to quantify departures from complete spatial randomness [43]. For each cell type, we computed *L*(*r*) − *r* curves at radii scaled by the median nearest-neighbor spacing. To test sex differences for microglia, we selected an *r* band using a pooled cluster-based one-sample test against the complete spatial randomness baseline, then computed a per-sample magnitude metric as

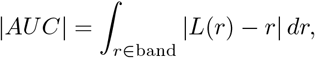

and evaluated male–female differences using a two-sided permutation test on the median difference. This procedure follows the exact implementation used to generate Fig. 5.

## 6 Declarations

### Ethics approval and consent to participate

Not applicable. All datasets used in this study were publicly available and de-identified.

### Consent for publication

Not applicable.

## Code availability

The `h5adify` source code is publicly available at: https://github.com/LOCImm/h5adify.git. All analyses presented in this study, including benchmarking, simulation experiments, and figure generation scripts, are available in a separate reproducibility repository at: https://github.com/LOCImm/h5adify_bench.git.

### Availability of data and materials

All benchmark datasets are publicly available from their respective repositories (for example, GEO and CZ CELLxGENE). The `h5adify` source code, benchmarking scripts, and curated gold-standard mappings are available in the associated public repository.

### Funding

This work was supported by the Agence Nationale de la Recherche (ANR) JCJC LOCImm (ANR-23-CE17-0027-01), and by the BRAINTWIN project funded under France 2030 through the PEPR Santé Numérique programme (ref. 2025-PEPR-121554). Additional support was provided by the MultiPOLA project funded by the Institut national du cancer (INCa; OSIRIS25), and by the CHANGING network of excellence against glioblastomas (INCa call LABREXCMP25; coordinator A.I.).

Spatial analyses and clinical translation were additionally supported by INCa and DGOS PRT-K projects, including Andro-iGlio (PRT-K23-096 and PRT-K24-096), and by the SiRIC CURAMUS programme (INCa-DGOS-Inserm 12560). T.K. acknowledges support from the European Union’s Horizon Europe research and innovation programme Cofund SOUND.AI under the Marie Sklodowska-Curie Grant Agreement No. 101081674.

### Authors’ contributions

L.R.R. and A.A. conceptualized the study. L.R.R. implemented the `h5adify` pipeline with input from A.M., M.N., and E.D. T.K. and S.G. contributed to the software engineering and integration benchmarks. A.I. and M.V. contributed clinical interpretation and study design for the glioblastoma analyses. K.L. and I.H.V. contributed to genomic and tumor microenvironment analyses and interpretation. L.R.R. and A.A. wrote the manuscript with input from all authors. All authors reviewed and approved the final manuscript.

## Acknowledgements

We thank the original study authors and the maintainers of public repositories for making single-cell and spatial transcriptomics datasets available.

## Appendix A Supplementary Methods

### A.1 Controlled simulation suite for metadata heterogeneity

To evaluate metadata harmonization under known ground truth, we implemented a simulation suite in `h5adify_benchmark_st_improved_v36.py` that generates three scenarios (SimA–SimC) and deliberately injects heterogeneous metadata encodings. The simulations were designed to reproduce frequent failure modes observed in public repositories: inconsistent column naming, partial missingness, free-text values that require semantic normalization, and gene naming schemes that differ across studies.

Across simulations we generate raw counts under a negative binomial model with per-gene dispersion parameter *θ*. Baseline gene means are cell-type specific, and we introduce structured unwanted variation via multiplicative perturbations of the mean, *µ*_*ij*_ ← *µ*_*ij*_ exp(*ε*_*j*_), where *ε*_*j*_ is sampled for a subset of genes and depends on the factor being simulated (batch, donor, or technology). This formulation yields realistic shifts that affect only a fraction of genes and preserves sparse count structure.

#### Gene naming heterogeneity and sex marker robustness

For each simulated dataset we generate a gene name vector whose format depends on the dataset index: (i) mixed symbols and synthetic names (e.g., `GENE00001`), (ii) lowercase variants, or (iii) Ensembl-like identifiers for non-special genes. Importantly, we always keep sex-marker genes as symbols (`XIST/Xist` and a panel of Y-linked genes) so that expression-based sex inference remains possible even when most genes are Ensembl-like.

##### A.1.1 SimA: Human brain single-cell simulation

SimA models a multi-study human brain single-cell setting with moderate confounding and inconsistent batch identifiers.

- Configuration: *n*_cells_ = 6000, *n*_genes_ = 6000, *n*_celltypes_ = 10, *n*_donors_ = 3, *θ* = 14.
- Effect structure: batch strength 1.5 applied to 30% of genes; donor strength 0.9 applied to 20% of genes; library-size strength 0.60.
- Studies: three simulated studies (`study1`, `study2`, `study3`) with batch labels corrupted to include variants such as `Study-02` and `S3`.

##### A.1.2 SimB: Glioblastoma-like human single-cell simulation

SimB models a glioblastoma-like scenario with stronger confounding, greater cell-type diversity, and technology variation across studies.

- Configuration: *n*_cells_ = 6500, *n*_genes_ = 7000, *n*_celltypes_ = 12, *n*_donors_ = 4, *θ* = 12.
- Effect structure: batch strength 1.8 applied to 35% of genes; donor strength 1.1 applied to 25% of genes; library-size strength 0.65.
- Studies: two studies (`gbm_like_1`, `gbm_like_2`) with technology labels `10xv2` versus `10xv3` and batch labels normalized to heterogeneous encodings such as `GBM-1` and `gbm2`.

##### A.1.3 SimC: Mouse Visium-like spatial simulation with multi-factor confounding

SimC models Visium-like spatial transcriptomics and explicitly combines section- and technology-level effects to mimic the more complex confounding structure typical of spatial cohorts.

- Configuration: *n*_cells_ = 6500 synthetic spots, *n*_genes_ = 6000, *n*_celltypes_ = 9, *n*_donors_ = 4, *θ* = 14.
- Effect structure: batch strength 1.35 applied to 25% of genes; donor strength 0.85 applied to 18% of genes; technology strength 2.6 applied to 45% of genes; library-size strength 0.70.
- Sections and technologies: three sections (`sec1`{`sec3`) crossed with three technologies (`Visium`, `Stereo-seq`, `SlideSeqV2`), generating nine datasets.
- Spatial patterns: each section draws a spatial pattern identifier from a predefined list that includes `cold`, `mixed`, `compartmentalized`, `addictive`, `exclusive`, `layer`, and gyrus. Pattern selection is deterministic given the section index, enabling reproducible benchmarks.

Spatial mixtures are generated with Spider [41]. For each cell type *k* Spider constructs a spatial score surface *s*_*k*_(*x, y*) according to the selected pattern regime. Probabilities are computed by a stabilized softmax,

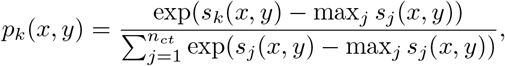

and spot-level mixtures are sampled from *p*_*k*_(*x, y*) before count generation. We then define a combined batch label `batch = section_technology` and store a `true_batch` string for evaluation.

##### A.1.4 Metadata corruption model

To emulate public metadata idiosyncrasies, we apply a corruption operator to each simulated dataset after generation. The operator performs the following actions (with scenario-specific probabilities):

- renames canonical columns to plausible alternatives (e.g., `donor` →`patient_id`, `batch` →`run` or `dataset`);
- injects additional batch-like columns (e.g., `lane`, `library`, `chemistry`) that partially correlate with the true batch;
- introduces missingness and conflicting values (e.g., partial sex labels, free-text disease strings);
- perturbs categorical formatting (case changes, punctuation, mixed delimiters) to require semantic and rule-based normalization.

The resulting pre-harmonization datasets therefore contain the correct information, but in encodings that would commonly break automated downstream workflows without explicit normalization.

## Appendix B Supplementary Notes on Reproducibility

All figures in this manuscript were generated from the scripts provided with the repository, with fixed random seeds and explicit configuration objects. The simulation benchmarking figure (Fig. 2) corresponds to the nine-panel report generated by the simulation suite and is additionally provided as a vector PDF to support inspection of individual scIB metrics.

---

**Fig. A1:**
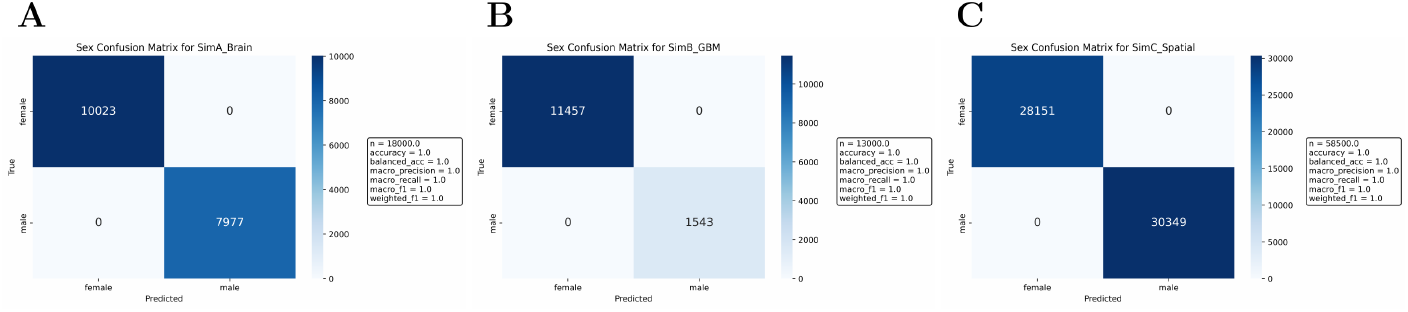
Supplementary confusion matrices for the sex prediction in the simulated datasets.

## References

[1] Virshup, I., Rybakov, S., Theis, F.J., Angerer, P., Wolf, F.A.: anndata: Annotated data. Journal of Open Source Software 9(101), 4371 (2024) 10.21105/joss.04371

[2] Wolf, F.A., Angerer, P., Theis, F.J.: SCANPY: large-scale single-cell gene expression data analysis. Genome Biology 19(1), 15 (2018) 10.1186/s13059-017-1382-0

[3] Palla, G., Spitzer, H., Klein, M., Fischer, D., Schaar, A.C., Kuemmerle, L.B., Rybakov, S., Ibarra, I.L., Holmberg, O., Virshup, I., Lotfollahi, M., Richter, S., Theis, F.J.: Squidpy: a scalable framework for spatial omics analysis. Nature Methods 19(2), 171–178 (2022) 10.1038/s41592-021-01358-2

[4] Program, C.C.S., Abdulla, S., Aevermann, B., Assis, P., Badajoz, S., Bell, S.M., Bezzi, E., Cakir, B., Chaffer, J., Chambers, S., Cherry, J., Chi, T., Chien, J., Dorman, L., Garcia-Nieto, P., Gloria, N., Hastie, M., Hegeman, D., Hilton, J., Huang, T., Infeld, A., Istrate, A.-M., Jelic, I., Katsuya, K., Kim, Y.J., Liang, K., Lin, M., Lombardo, M., Marshall, B., Martin, B., McDade, F., Megill, C., Patel, N., Predeus, A., Raymor, B., Robatmili, B., Rogers, D., Rutherford, E., Sadgat, D., Shin, A., Small, C., Smith, T., Sridharan, P., Tarashansky, A., Tavares, N., Thomas, H., Tolopko, A., Urisko, M., Yan, J., Yeretssian, G., Zamanian, J., Mani, A., Cool, J., Carr, A.: Cz cellxgene discover: a single-cell data platform for scalable exploration, analysis and modeling of aggregated data. Nucleic Acids Research 53(D1), 886–900 (2024) 10.1093/nar/gkae1142 https://academic.oup.com/nar/article-pdf/53/D1/D886/60882786/gkae1142.pdf

[5] Theodoris, C.V., Xiao, L., Chopra, A., Chaffin, M.D., et al.: Transfer learning enables predictions in network biology. Nature 618(7965), 616–624 (2023) 10.1038/s41586-023-06139-9

[6] Cui, H., Wang, C., Maan, H., et al.: scgpt: toward building a foundation model for single-cell multi-omics using generative ai. Nature Methods 21(8), 1470–1480 (2024) 10.1038/s41592-024-02201-0

[7] Hao, M., Gong, J., Zeng, X., Liu, C., et al.: Large-scale foundation model on single-cell transcriptomics. Nature Methods 21(8), 1481–1491 (2024) 10.1038/s41592-024-02305-7

[8] Heumos, L., Schaar, A.C., Lance, C., et al.: Best practices for single-cell analysis across modalities. Nature Reviews Genetics 24(8), 550–572 (2023) 10.1038/s41576-023-00586-w

[9] Bernstein, M.N., Doan, A., Dewey, C.N.: Metasra: normalized human sample-specific metadata for the sequence read archive. Bioinformatics 33(18), 2914–2923 (2017) 10.1093/bioinformatics/btx334

[10] Wilks, C., Zheng, S.C., Chen, F.Y., Charles, R., Solomon, B., Ling, J.P., et al.: recount3: summaries and queries for large-scale rna-seq expression and splicing. Genome Biology 22(1), 323 (2021) 10.1186/s13059-021-02533-6

[11] Li, Z., Li, J., Yu, P.: Geometacuration: a web-based application for accurate manual curation of gene expression omnibus metadata. Database (Oxford), 019 (2018) 10.1093/database/bay019

[12] Lopez, R., Regier, J., Cole, M.B., Jordan, M.I., Yosef, N.: Deep generative modeling for single-cell transcriptomics. Nature Methods 15(12), 1053–1058 (2018) 10.1038/s41592-018-0229-2

[13] Gayoso, A., Lopez, R., Xing, G., Boyeau, P., Valiollah Pour Amiri, V., Hong, J., et al.: A python library for probabilistic analysis of single-cell omics data. Nature Biotechnology 40(2), 163–166 (2022) 10.1038/s41587-021-01206-w

[14] Stuart, T., Butler, A., Hoffman, P., Hafemeister, C., Papalexi, E., Mauck, W.M., et al.: Comprehensive integration of single-cell data. Cell 177(7), 1888–190221 (2019) 10.1016/j.cell.2019.05.031

[15] Korsunsky, I., Millard, N., Fan, J., Slowikowski, K., Zhang, F., Wei, K., et al.: Fast, sensitive and accurate integration of single-cell data with harmony. Nature Methods 16(12), 1289–1296 (2019) 10.1038/s41592-019-0619-0

[16] Hie, B., Bryson, B., Berger, B.: Efficient integration of heterogeneous single-cell transcriptomes using scanorama. Nature Biotechnology 37(6), 685–691 (2019) 10.1038/s41587-019-0113-3

[17] Tasic, B., Yao, Z., Graybuck, L.T., Smith, K.A., Nguyen, T.N., Bertagnolli, D., Goldy, J., Garren, E., Economo, M.N., Viswanathan, S., Penn, O., Bakken, T., Menon, V., Miller, J., Fong, O., Hirokawa, K.E., Lathia, K., Rimorin, C., Tieu, M., Larsen, R., Casper, T., Barkan, E., Kroll, M., Parry, S., Shapovalova, N.V., Hirschstein, D., Pendergraft, J., Sullivan, H.A., Kim, T.K., Szafer, A., Dee, N., Groblewski, P., Wickersham, I., Cetin, A., Harris, J.A., Levi, B.P., Sunkin, S.M., Madisen, L., Daigle, T.L., Looger, L., Bernard, A., Phillips, J., Lein, E., Hawrylycz, M., Svoboda, K., Jones, A.R., Koch, C., Zeng, H.: Shared and distinct transcriptomic cell types across neocortical areas. Nature 563(7729), 72–78 (2018) 10.1038/s41586-018-0654-5

[18] Han, X., Zhou, Z., Fei, L., Sun, H., Wang, R., Chen, Y., Chen, H., Wang, J., Tang, H., Ge, W., Zhou, Y., Ye, F., Jiang, M., Wu, J., Xiao, Y., Jia, X., Zhang, T., Ma, X., Zhang, Q., Bai, X., Lai, S., Yu, C., Zhu, L., Lin, R., Gao, Y., Wang, M., Wu, Y., Zhang, J., Zhan, R., Zhu, S., Hu, H., Wang, C., Chen, M., Huang, H., Liang, T., Chen, J., Wang, W., Zhang, D., Guo, G.: Construction of a human cell landscape at single-cell level. Nature 581(7808), 303–309 (2020) 10.1038/s41586-020-2157-4

[19] Almanzar, N., Antony, J., Baghel, A.S., Bakerman, I., Bansal, I., Barres, B.A., Beachy, P.A., Berdnik, D., Bilen, B., Brownfield, D., Cain, C., Chan, C.K.F., Chen, M.B., Clarke, M.F., Conley, S.D., Darmanis, S., Demers, A., Demir, K., Morree, A., Divita, T., Bois, H., Ebadi, H., Espinoza, F.H., Fish, M., Gan, Q., George, B.M., Gillich, A., Gòmez-Sjöberg, R., Green, F., Genetiano, G., Gu, X., Gulati, G.S., Hahn, O., Haney, M.S., Hang, Y., Harris, L., He, M., Hosseinzadeh, S., Huang, A., Huang, K.C., Iram, T., Isobe, T., Ives, F., Jones, R.C., Kao, K.S., Karkanias, J., Karnam, G., Keller, A., Kershner, A.M., Khoury, N., Kim, S.K., Kiss, B.M., Kong, W., Krasnow, M.A., Kumar, M.E., Kuo, C.S., Lam, J., Lee, D.P., Lee, S.E., Lehallier, B., Leventhal, O., Li, G., Li, Q., Liu, L., Lo, A., Lu, W.-J., Lugo-Fagundo, M.F., Manjunath, A., May, A.P., Maynard, A., McGeever, A., McKay, M., McNerney, M.W., Merrill, B., Metzger, R.J., Mignardi, M., Min, D., Nabhan, A.N., Neff, N.F., Ng, K.M., Nguyen, P.K., Noh, J., Nusse, R., Pálovics, R., Patkar, R., Peng, W.C., Penland, L., Pisco, A.O., Pollard, K., Puccinelli, R., Qi, Z., Quake, S.R., Rando, T.A., Rulifson, E.J., Schaum, N., Segal, J.M., Sikandar, S.S., Sinha, R., Sit, R.V., Sonnenburg, J., Staehli, D., Szade, K., Tan, M., Tan, W., Tato, C., Tellez, K., Dulgeroff, L.B.T., Travaglini, K.J., Tropini, C., Tsui, M., Waldburger, L., Wang, B.M., Weele, L.J., Weinberg, K., Weissman, I.L., Wosczyna, M.N., Wu, S.M., Wyss-Coray, T., Xiang, J., Xue, S., Yamauchi, K.A., Yang, A.C., Yerra, L.P., Youngyunpipatkul, J., Yu, B., Zanini, F., Zardeneta, M.E., Zee, A., Zhao, C., Zhang, F., Zhang, H., Zhang, M.J., Zhou, L., Zou, J., The Tabula Muris Consortium: A single-cell transcriptomic atlas characterizes ageing tissues in the mouse. Nature 583(7817), 590–595 (2020) 10.1038/s41586-020-2496-1

[20] Travaglini, K.J., Nabhan, A.N., Penland, L., Sinha, R., Gillich, A., Sit, R.V., Chang, S., Conley, S.D., Mori, Y., Seita, J., Berry, G.J., Shrager, J.B., Metzger, R.J., Kuo, C.S., Neff, N., Weissman, I.L., Quake, S.R., Krasnow, M.A.: A molecular cell atlas of the human lung from single-cell RNA sequencing. Nature 587(7835), 619–625 (2020) 10.1038/s41586-020-2922-4

[21] He, S., Jin, Y., Nazaret, A., Shi, Y., Chen, R., et al.: Starfysh integrates spatial transcriptomic and histologic data to reveal heterogeneous tumor–immune hubs. Nature Biotechnology 43(2), 223–235 (2025) 10.1038/s41587-024-02173-8

[22] Ravi, V.M., Will, P., Kueckelhaus, J., Sun, N., Joseph, K., Salié, H., et al.: Spatially resolved multi-omics deciphers bidirectional tumor-host interdependence in glioblastoma. Cancer Cell 40(6), 639–65513 (2022) 10.1016/j.ccell.2022.05.009

[23] Luecken, M.D., Büttner, M., Chaichoompu, K., et al.: Benchmarking atlas-level data integration in single-cell genomics. Nature Methods 19(1), 41–50 (2022) 10.1038/s41592-021-01336-8

[24] Mondal, R., Sen, M., Sengupta, S., Maity, W., Palapetta, S., Dhruw, N.K., Jha, A.: Multi-agent ai system for high quality metadata curation at scale. bioRxiv (2025) 10.1101/2025.06.10.658658 https://www.biorxiv.org/content/early/2025/06/11/2025.06.10.658658.full.pdf

[25] Verbitsky, A., Boutet, P., Eslami, M.: Metadata harmonization from biological datasets with language models. Bioinformatics Advances 5(1), 241 (2025) 10.1093/bioadv/vbaf241 https://academic.oup.com/bioinformaticsadvances/article-pdf/5/1/vbaf241/64457803/vbaf241.pdf

[26] Nouri, N., Artzi, R., Savova, V.: An agentic AI framework for ingestion and standardization of single-cell RNA-seq data analysis. npj Artificial Intelligence 2(1), 8 (2026) 10.1038/s44387-025-00064-0

[27] Schaefer, M., Peneder, P., Malzl, D., Lombardo, S.D., Peycheva, M., Burton, J., Hakobyan, A., Sharma, V., Krausgruber, T., Sin, C., Menche, J., Tomazou, E.M., Bock, C.: Multimodal learning enables chat-based exploration of single-cell data. Nature Biotechnology (2025) 10.1038/s41587-025-02857-9

[28] Wu, Y., Tang, F.: scextract: leveraging large language models for fully automated single-cell RNA-seq data annotation and prior-informed multi-dataset integration. Genome Biology 26(1), 174 (2025) 10.1186/s13059-025-03639-x

[29] Gao, R., Bai, S., Henderson, Y.C., Lin, Y., Schalck, A., Yan, Y., et al.: Delineating copy number and clonal substructure in human tumors from single-cell transcriptomes. Nature Biotechnology 39(5), 599–608 (2021) 10.1038/s41587-020-00795-2

[30] Mi, X., Ye, Z.-L., Zhang, X., Chen, X., Dai, X., et al.: Sex- and age-differences in the expression of critical blood-brain barrier regulators: a physiological context. Biology of Sex Differences 16(1), 67 (2025) 10.1186/s13293-025-00751-2

[31] Wälchli, T., Ghobrial, M., Schwab, M., Takada, S., et al.: Single-cell atlas of the human brain vasculature across development, adulthood and disease. Nature 632(8025), 603–613 (2024) 10.1038/s41586-024-07493-y

[32] Efremova, M., Vento-Tormo, R., Teichmann, S.A., Vento-Tormo, R.: Cellphonedb: inferring cell–cell communication from combined expression of multi-subunit ligand–receptor complexes. Nature Protocols 15(4), 1484–1506 (2020) 10.1038/s41596-020-0292-x

[33] Jin, S., Guerrero-Juarez, C.F., Zhang, L., et al.: Inference and analysis of cell-cell communication using cellchat. Nature Communications 12(1), 1088 (2021) 10.1038/s41467-021-21246-9

[34] Browaeys, R., Saelens, W., Saeys, Y.: Nichenet: modeling intercellular communication by linking ligands to target genes. Nature Methods 17(2), 159–162 (2020) 10.1038/s41592-019-0667-5

[35] Armingol, E., Baghdassarian, H.M., Lewis, N.E., et al.: The diversification of methods for studying cell-cell interactions and communication. Nature Reviews Genetics 25(6), 381–400 (2024) 10.1038/s41576-023-00685-8

[36] Bayik, D., Zhou, Y., Park, C., Hong, C., Vail, D., Silver, D.J., et al.: Myeloid-derived suppressor cell subsets drive glioblastoma growth in a sex-specific manner. Cancer Discovery 10(8), 1210–1225 (2020) 10.1158/2159-8290.CD-19-1355

[37] Ochocka, N., Segit, P., Walentynowicz, K.A., Wojnicki, K., Cyranowski, S., Swatler, J., et al.: Single-cell rna sequencing reveals functional heterogeneity of glioma-associated brain macrophages. Nature Communications 12(1), 1151 (2021) 10.1038/s41467-021-21407-w

[38] Ochocka, N., Segit, P., Wojnicki, K., Cyranowski, S., et al.: Specialized functions and sexual dimorphism explain the functional diversity of the myeloid populations during glioma progression. Cell Reports 42(1), 111971 (2023) 10.1016/j.celrep.2022.111971

[39] Hiller-Vallina, S., Mondejar-Ruescas, L., Caamaño-Moreno, M., et al.: Sexual-biased necroinflammation is revealed as a predictor of bevacizumab benefit in glioblastoma. Neuro-Oncology 26(7), 1213–1227 (2024) 10.1093/neuonc/noae033

[40] Greenwald, A.C., Darnell, N.G., Hoefflin, R., Simkin, D., Mount, C.W., Gonzalez Castro, L.N., et al.: Integrative spatial analysis reveals a multi-layered organization of glioblastoma. Cell 187(10), 2485–250126 (2024) 10.1016/j.cell.2024.03.029

[41] Yang, J., Wei, N., Qu, Y., Wu, H.-J., Zheng, X., et al.: Spider: a flexible and unified framework for simulating spatial transcriptomics data. Bioinformatics 42(1), 562 (2026) 10.1093/bioinformatics/btaf562

[42] Johnson, W.E., Li, C., Rabinovic, A.: Adjusting batch effects in microarray expression data using empirical bayes methods. Biostatistics 8(1), 118–127 (2007) 10.1093/biostatistics/kxj037

[43] Ripley, B.D.: The second-order analysis of stationary point processes. Journal of Applied Probability 13(2), 255–266 (1976) 10.2307/3212829

